# Supramolecular Biomimetic Topology: Macrophage-Engaged Neoadjuvant Platform for Preoperative Immunotherapy

**DOI:** 10.1101/2025.03.02.641066

**Authors:** Huayu Wu, Dan Zhong, Zhongwei Gu

## Abstract

The persistent immunosuppression and spatial heterogeneity of advanced solid tumors necessitate next-generation neoadjuvant platforms capable of reprogramming tumor microenvironments while overcoming resistance to immune checkpoint blockade. Addressing this unmet clinical challenge, we report a β-glucan-empowered nano-neoadjuvant agents (NNAs) engineered through topology-guided supramolecular assembly of pathogen-mimetic β-glucan motifs onto photoresponsive single-wall carbon nanotubes. This biomimetic architecture capitalizes on three synergistic mechanisms: (1) Dectin-1-mediated pathogen-associated molecular pattern recognition to rewire myeloid cell functionality, (2) endogenous leukocyte-hitchhiking delivery paradigm, and (3) spatiotemporally controlled photothermal ablation via near-infrared transducing SWNT cores. NNAs also orchestrates a proinflammatory cascade involving IL-12-dependent Th1 polarization and activation, effectively converting “cold” tumors into immunogenic niches. This strategy redefines preoperative therapeutic paradigms, offering a multimodal platform to harness the proinflammatory adjuvant effect of SWNTs for preoperative immunomodulation in advanced-stage malignancies. By bridging innate immune recognition with nanotechnology-enabled precision, this paradigm establishes a new roadmap for preoperative conditioning of advanced malignancies, offering a theranostic framework to transform surgical outcomes through immunologically informed nanomedicine design.

## Introduction

The management of advanced solid cancers is a multidisciplinary approach, where locoregional therapies such as surgery or thermal therapy, along with systemic therapy, have become the standard-of-care modalities ^1-3^. The neoadjuvant immune approaches, such as immune checkpoint inhibitors (ICIs), have emerged as promising post-surgical treatment modality for eliciting systemic anti-tumor responses against micrometastatic disease ^4-6^. However, pathological responsiveness of ICI is highly limited by the immunosuppressive microenvironment and the heterogeneous intratumoral components, particularly macrophages and monocytes which constitute the majority of infiltrating immune cells ^7-10^. Regarding the heterogeneous resistance of neoadjuvant ICI therapies among patients, it is crucial to develop integrated nano-based neoadjuvant immune approaches to establish systemic anti-tumor immunity establishment prior to tumor resection.

Biological molecules such as polypeptide and polysaccharides may function as allergens or pathogen-associated molecular patterns (PAMPs) ^11-13^. β-glucans, are polysaccharides constructed of glucose monomers with backbone of linear β-(1→3)-D-glucans with branched β-(1→6)-D-glucose side chains, which interact with C-type lectin receptor (dectin-1) on myeloid immune cells, such as macrophages and monocytes, for innate immune activating ^14, 15^. The interactions with dectin-1 depend highly on the morphology of particles, thereby regulating the adjuvant effects that activate inflammatory innate immune responses, promote cytokine production, and elicit adaptive B/T cell activation ^16-19^. β-glucans form helical complexes by triple-stranded macromolecular in natural state. In this work, we promoted a supramolecular hybrid structure by self-assembling of the β-glucans on single wall carbon nanotubes (SWNTs) as an innovative nano-agents for alternative neoadjuvant treatment.

Nanostructured particles hold inherent interactions with phagocytic myeloid cells, and are thus ideal tools for immunomodulation ^20, 21^. And these interactions can be versatilely controlled by shape, size, properties of surface properties, and compositions of nano-assemblies ^22-26^. Through rational design of the bioactive nano-assemblies, it might allow multimodal immune eliciting as revolutionary neoadjuvant immune-regulating platforms. Control over surface activity and morphology may also allow for enhanced target specificity and designed immune eliciting. SWNTs have a broad range of potential in applications of drug delivery and photothermal therapy ^27^. Meanwhile, the innate interaction of SWNTs with myeloid immune cells can amplify the proinflammatory response as adjuvants, which are highly well-controlled when SWMTs are engineered to carry bio-active molecules modulating the activity of systemic immunity ^28-33^. To leverage the intrinsic immunomodulatory properties of SWNTs, we engineered a β-glucan-functionalized nanoarchitecture through rational polysaccharide design. This strategic functionalization enables precise spatiotemporal control over immune activation dynamics, directly addressing the unmet need for preoperative immunomodulatory precision in advanced solid tumor therapy. And the efficient hyperpyrexia therapy upon NIR irradiation at tumor site can also triggers immunological responses for multimodal therapy ^34^. In this work, through surface modified by the bio-active β-glucan polysaccharides, the accurate assemblies of nanoscopic hybrids, which we defined as nano-neoadjuvant agents (NNAs), were supposed to modulating the systemic immune responses and avoid undesirable effects such as cytotoxicity.

In the hybrid structure, SWNTs are high-aspect-ratio templates as rigid cores for construction of nano-assemblies. β-glucans enwind on SWNTs by helix-forming through the hydrophobic side of glucose monomers interacting with the superhydrophobic nanotube walls. The bioactive surface modification and elongated morphology endow the NNAs with efficient activation of dectin-1 on myeloid immune cells and systemic immunity boosting. Meanwhile, the activated myeloid immune cells hitchhiked by NNAs also show efficient tumor tropism guided by chemokine gradient, and thus cause superior photothermal therapy for locoregional and systemic control as a novel nano-neoadjuvant treatment post-surgery.

## Results and discussion

### Preparation and characterization of NNAs

An oxidizing cutting process utilized to yield short SWNTs as chemical scissorsis. The first step requires the breakage of carbon-carbon bonds in the lattice while the second step is aimed at etching at these damage sites to create short, cut nanotubes. Then, co-dissolution of SWNTs and β-glucans and in a co-solvent system (e.g., DMSO) facilitates supramolecular self-assembly upon aqueous phase transition, yielding monodisperse core-shell nanostructures through synergistic CH-π interactions and hydrophilic patterning ^35^. The formation of NNAs was mediated by hydrophobic interactions between β-glucan chains and SWNTs. This molecular rearrangement enabled the encapsulation of SWNTs within the central cavity of the polysaccharide’s helical architecture. The black NNAs suspension exhibited stable dispersion in aqueous solution. TEM analysis (Figure 1A) revealed a uniform β-glucan polysaccharide coating on the surface of shortened SWNTs, whereas the non-cut SWNTs exhibited pronounced aggregation due to insufficient surface modification (Supplementary Fig. 1 and Fig. 2). CD spectroscopic measurements (Figure 1B) also demonstrated the successful composition of unique nano-structure of NNAs. UV-vis spectroscopic measurements (Figure 1C, left) and raman measurements (Figure 1C, right) characteristic spectral lines confirmed the core-shell nanostructures in which the SWNTs were entrapped within the helical superstructure of β-1,3-glucans.

**Figure 1.**
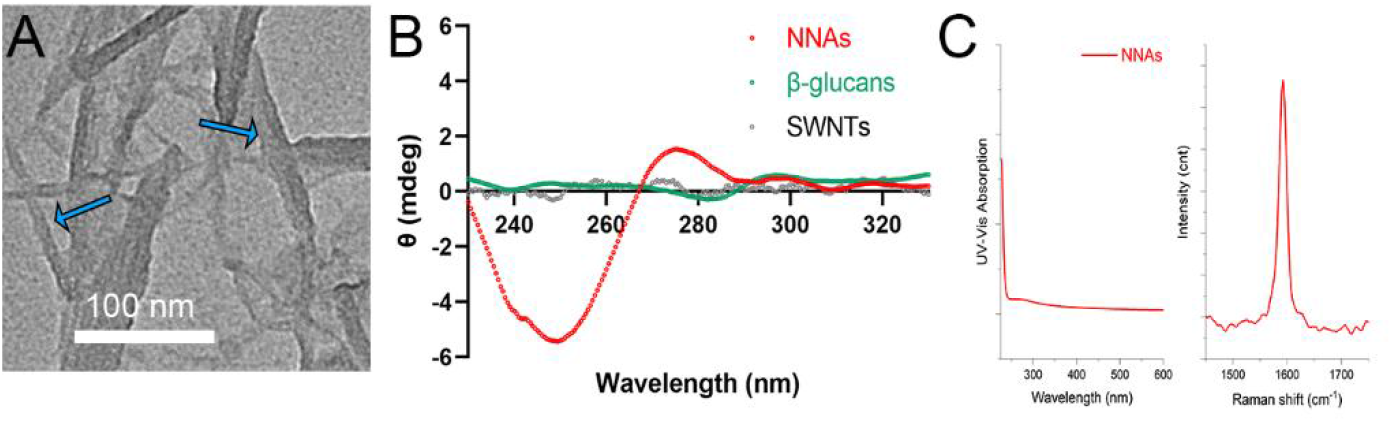
Preparation and characterization of NNAs. (A) TEM images of NNAs, (B) CD spectroscopic measurements, (C) UV-vis spectroscopic and raman measurements.

**Figure 2.**
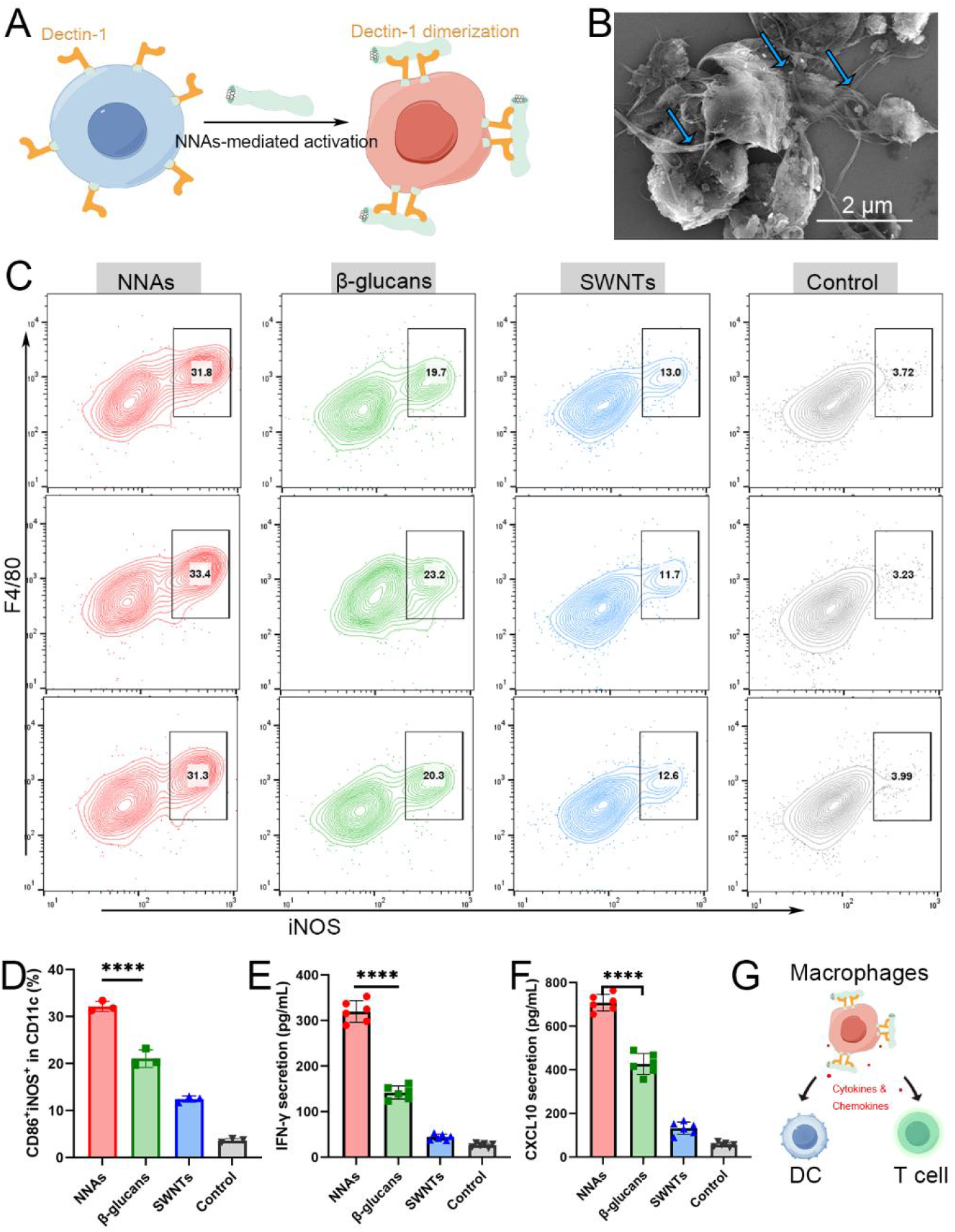
Nano-bio interactions of NNAs with macrophages. (A) NNAs-mediated dectin-1 clustering and macrophage activation. (B) SEM reveals NNAs (blue arrows) binding to macrophage membranes. (C) flow cytometry results and qualifications showed M1-polarizing BMDMs (F4/80^+^iNOS^+^) functional dominance. (E) IFN-γ concentration and (F) CXCL10 concentration in the supernates of BMDMs treated with the formulations. (G) macrophage-centric activation cascade.

The rod-like NNAs are constructed by anchoring functionalized β-1,3-glucans onto the surface of single-walled carbon nanotubes (SWNTs), forming a high-aspect-ratio architecture. This unique surface functionalization leverages the bioactive properties of β-1,3-glucans, which are distinctively absent in conventional polysaccharides. The synergistic combination of the anisotropic morphology and surface-exposed polysaccharide moieties significantly enhances nano-bio interactions in vitro, particularly improving the immunomodulatory efficacy of the biomimetic neoadjuvant platform through optimized engagement with immune cells during preoperative conditioning.

These multimodal datasets collectively establish that the high-aspect-ratio morphology synergizes with surface-exposed β-glucans to emulate pathogen-associated molecular patterns, thereby amplifying immune surveillance pathways essential for preoperative neoadjuvant applications.

### Nano-bio interactions *in vitro*

The bio-behavior of the polysaccharides-modified hybrid of nano-assemblies, can be effectively adjusted to control nano-bio interactions. In particular, cell-uptake efficiency of the resultant polysaccharide/polynucleotide complexes was remarkably enhanced when functional groups recognized in a biological system were introduced as pendent groups.

The immunostimulatory potency of β-glucans-functionalized NNAs was rigorously validated through a multidisciplinary investigation of immune cell responses. The rod-shaped NNAs with bioactive polysaccharide coatings may potently activate macrophages via dectin-1 clustering, the classical β-glucan receptor (Figure 2A) and morphology-driven phagocytosis. This dual activation mechanism amplifies pro-inflammatory responses in macrophages while bridging innate-adaptive immunity.

Macrophage interactions were dissected through combined SEM and confocal microscopy. SEM imaging of NNAs interacting with bone marrow-derived macrophages (BMDM) surfaces revealed ligand-receptor binding patterns characterized by localized membrane deformations (Figure 2B). Functional polarization assays revealed that NNAs induced M1-associated iNOS expression (Figure 2C) by 1.53-fold compared to soluble β-glucan group and 8.82-fold compared to untreated controls (Figure 2D). The expression of pro-inflammatory cytokine IFN-γ (Figure 2E) and Th1-associated chemokine CXCL10 (Figure 2F) also showed a marked increase, indicating the potent activation effect of NNAs on the macrophage system. Notably, this macrophage-centric activation cascade (Figure 2G) was further amplified by NNAs through enhanced dendritic cell (DC) maturation, as evidenced by elevated CD80/86 expression, ultimately priming a robust Th1-skewed T cell response.

Then, bone marrow-derived dendritic cells (BMDCs) exposed to NNAs (10 μg/mL, 24 h) exhibited significantly enhanced activation profiles. Flow cytometry analysis indicated that 44.4 ± 0.51% of BMDCs expressed both CD86 and CD80 co-stimulatory markers (Figure 3A, Figure 3B), a dramatic increase compared to the group treated with soluble β-glucan controls (p < 0.001). This significant upregulation of dual co-stimulatory marker expression (CD86^+^CD80^+^) strongly suggests that NNAs, as a nano-engineering strategy for polysaccharides, profoundly enhance DC activation through synergistic mechanisms. These nanoscale advantages collectively drive robust DC maturation, positioning NNAs as superior immunopotentiators for next-generation vaccine adjuvants.

**Figure 3.**
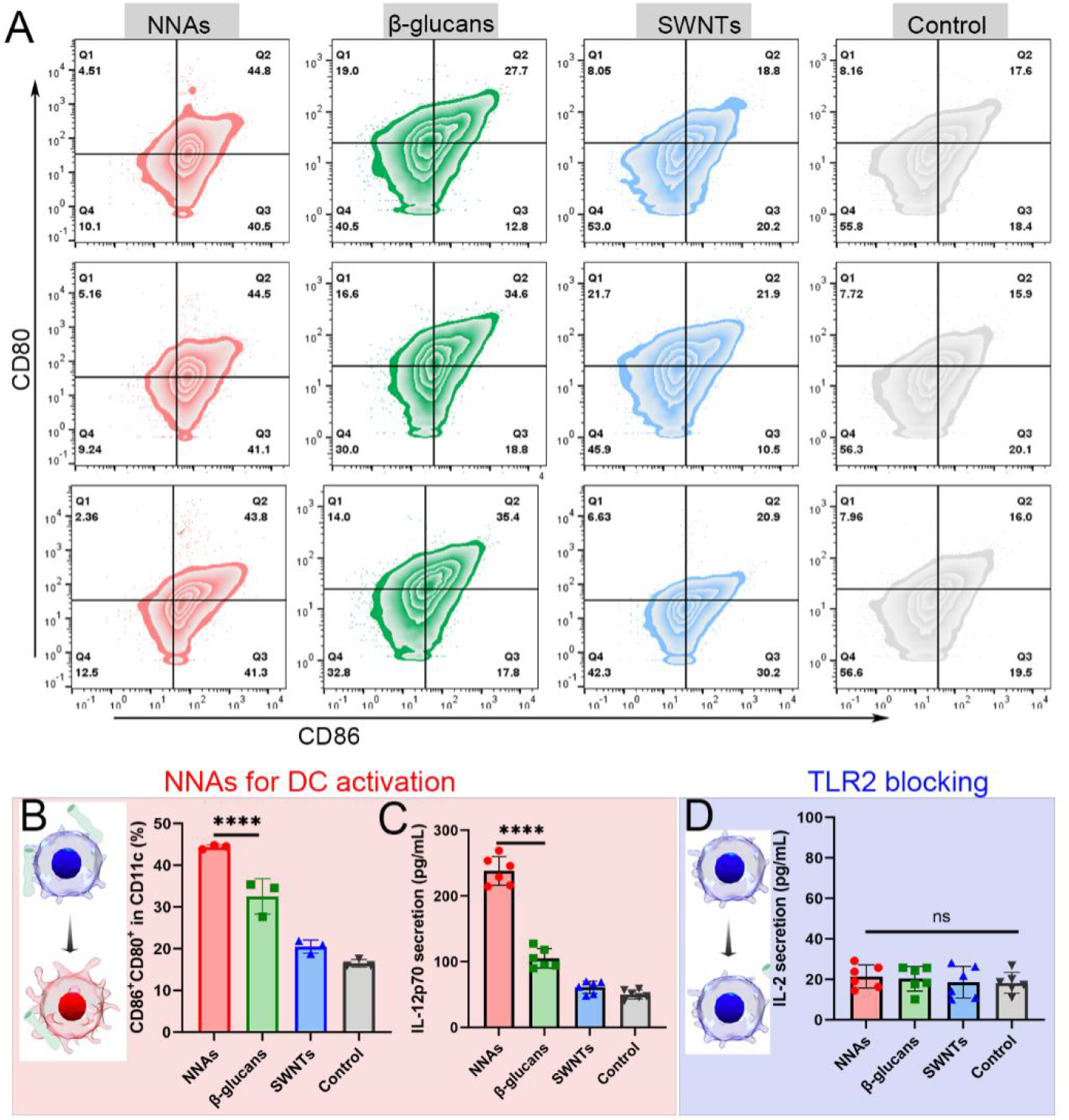
NNAs enhance dendritic cell activation. (A) Representative flow cytometry plots showing CD86 & CD80 co-expression in BMDCs treated with NNAs (10 μg/mL, 24 h), (B) Quantification of dual-positive (CD86^+^CD80^+^) cells, highlighting nanoscale-driven DC maturation, (C) IL-12p70 secretion by ELISA, (D) TAK-242 inhibition (1 μM) reducing IL-12 production by NNAs, (statistical analysis by one-way ANOVA, ****p < 0.001).

This activation was corroborated by ELISA quantification of IL-12p70 secretion, which reached 238.1 ± 19.9 pg/mL (Figure 3C), representing a 4.75-fold elevation over baseline levels. Pre-treatment with the TLR2-specific inhibitor TAK-242 (1 μM) reduced IL-12 production by NNAs (Figure 3D), confirming the critical role of β-glucan-mediated TLR2 engagement in dendritic cell activation.

T cell priming assays in coculture systems revealed cascading immunogenicity. DCs pre-treated with NNAs induced 28.1 ± 6.5% IFN-γ+ CD8+ T cells (Figure 4A and Supplementary Fig. 3), signifying enhanced effector function for immune killing.

**Figure 4.**
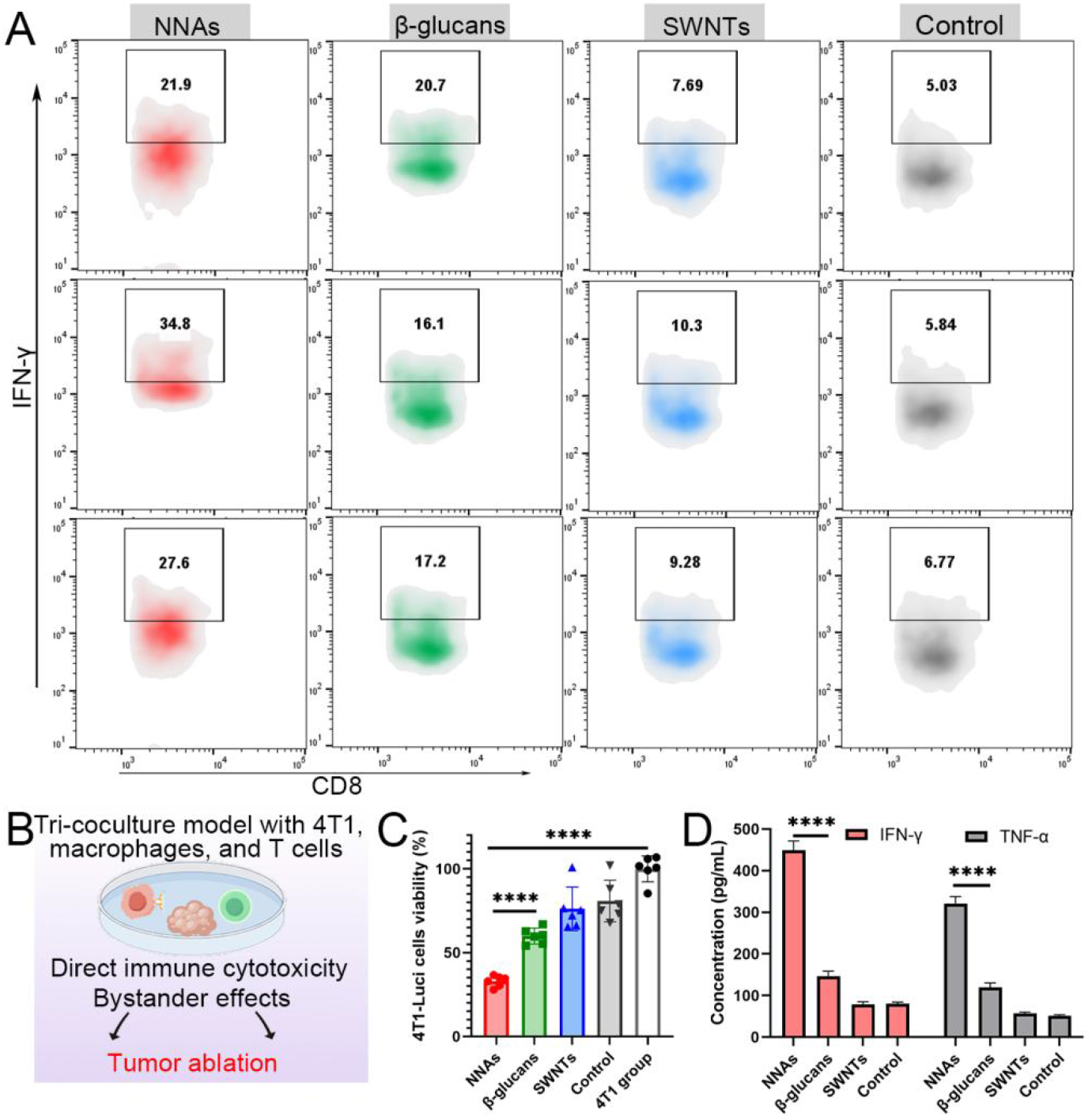
Multi-mechanism synergistic tumor ablation. (A) NNA-Pretreated Dendritic Cells Prime Antigen-Specific IFN-γ^+^ CD8^+^ T Cells. (B) Tri-Cell Coculture Model Recapitulates Tumor-Immune Microenvironment 4T1-Luc cells (tumor), M1-polarized macrophages (innate immunity), and activated CD8^+^ T cells (adaptive immunity) co-cultured at a 2:1:1 ratio. (C) Synergistic Tumor Killing in Tri-Coculture System NNA treatment reduced 4T1-Luc viability, outperforming free β-glucan. Data normalized to luciferase activity (RLU). (D) ELISA quantification of bystander cytokines in co-culture supernatants drives tumor ablation.

This study establishes a biomimetic nanotechnology platform that harnesses pathogen-mimicking structural and molecular cues to rewire immune recognition pathways. By integrating high-aspect-ratio nanostructures with surface-exposed β-glucans, the NNAs exploit evolutionary conserved mechanisms of microbial detection, effectively bridging innate and adaptive immune activation. The observed synergy between geometric cues and biochemical signaling highlights a fundamental design principle: nanostructure morphology dictates cellular interaction dynamics, while surface chemistry fine-tunes receptor-specific responses. Such dual modulation enables precise amplification of pro-inflammatory cascades without overt toxicity—a critical advancement over conventional adjuvants burdened by off-target effects. The demonstrated capacity to reverse immunosuppressive checkpoints and enhance phagocytic clearance underscores the potential of NNAs as a neoadjuvant strategy to precondition tumor microenvironments prior to surgery. Future efforts should address in vivo biodistribution challenges while exploring combinatorial regimens with checkpoint inhibitors, ultimately translating this pathogen-inspired nanotechnology into actionable immunotherapies *in vivo*.

These findings collectively position NNAs as a programmable neoadjuvant platform capable of reconditioning immunosuppressive niches through geometry-guided immune targeting and β-glucan-driven danger signaling—a strategy poised to enhance preoperative immunotherapy paradigms. The NNAs’ immunomodulatory supremacy stems from their biomimetic duality: the high-aspect-ratio architecture emulates bacterial morphology to exploit evolutionary conserved phagocytic mechanisms, while surface-anchored β-glucans serve as pathogen-associated molecular patterns (PAMPs) for targeted receptor activation. T This design outperforms conventional polysaccharide composites by circumventing helical entrapment that masks ligand accessibility. Notably, the observed phagocytic efficiency approaches levels induced by bacterial-like immune agonists (e.g., LPS or CpG motifs), thereby offering pioneering insights for biomimetic nanomedicine in neoadjuvant clinical applications.

### NNAs reprogram tumor-immune interactions in multicellular co-cultures

In a tri-culture model combining 4T1 breast cancer cells (luciferase-expressing), macrophages, and CD8+ T cells, NNAs treatment (10 μg/mL) is predicted to reduce 4T1 survival to approximately 33.0% (Figure 4B) compared to untreated controls (80.8% survival), demonstrating potent tumor-killing synergy between macrophages and T cells. Cytokine levels in the supernatant are expected to increase significantly as bystander effects for tumor killing (Figure 4C). IFN-γ concentrations reached 450 pg/mL (vs. 80 pg/mL in controls) and TNF-α 320 pg/mL (vs. 50 pg/mL), indicating robust immune activation. Notably, free β-glucan will likely show weaker effects (60% tumor survival, IFN-γ 150 pg/mL), highlighting the structural advantage of NNAs in enhancing immune cell coordination. Even when physically separated from tumor cells in transwell systems, NNAs achieved 53% tumor suppression (Supplementary Fig. 4), driven by soluble immune signals rather than direct contact. This bystander effect highlights their ability to amplify antitumor immunity beyond localized interactions.

Such topology-engineered NNAs further mimic pathogen-associated molecular patterns (PAMPs), surpassing conventional “stealth” nanocarriers by intentionally provoking controlled pro-inflammatory cascades, a strategy critical for reversing immunosuppressive tumor niches. The combined tumor-killing and immune-priming capabilities position NNAs as a promising neoadjuvant platform. By shrinking tumors pre-surgery and establishing “immune memory” against residual cancer cells, NNAs may following address two critical challenges in cancer surgery: reducing resection complexity and preventing postoperative recurrence.

### Tumor-Targeted Accumulation and Photothermal Ablation: NNAs as a Neoadjuvant Platform

The rod-like morphology of NNAs with high aspect ratio and their surface polysaccharide decoration (e.g., β-glucan ligands) confer exceptional macrophage tropism. Upon intravenous injection, NNAs rapidly bind to circulating macrophages via dectin-1 receptor interactions, leveraging macrophage-mediated chemotaxis to home to tumor lesions. This macrophage-driven tumor-targeting mechanism is particularly critical for neoadjuvant immunotherapy, where localized delivery of immune activators prior to surgery amplifies antitumor responses while minimizing systemic toxicity. To validate the preferential accumulation of NNAs in tumors, 4T1-bearing BALB/c mice were intravenously injected with NNAs. In vivo photoacoustic imaging revealed rapid NNA enrichment at tumor sites, peaking at 6 h post-injection (Figure 5A). This targeting efficiency was much higher than SWNTs, attributable to NNAs’ rod-shaped geometry and topologically covered appearance of polysaccharides for enhancing nano-bio interaction and tumor retention.

**Figure 5.**
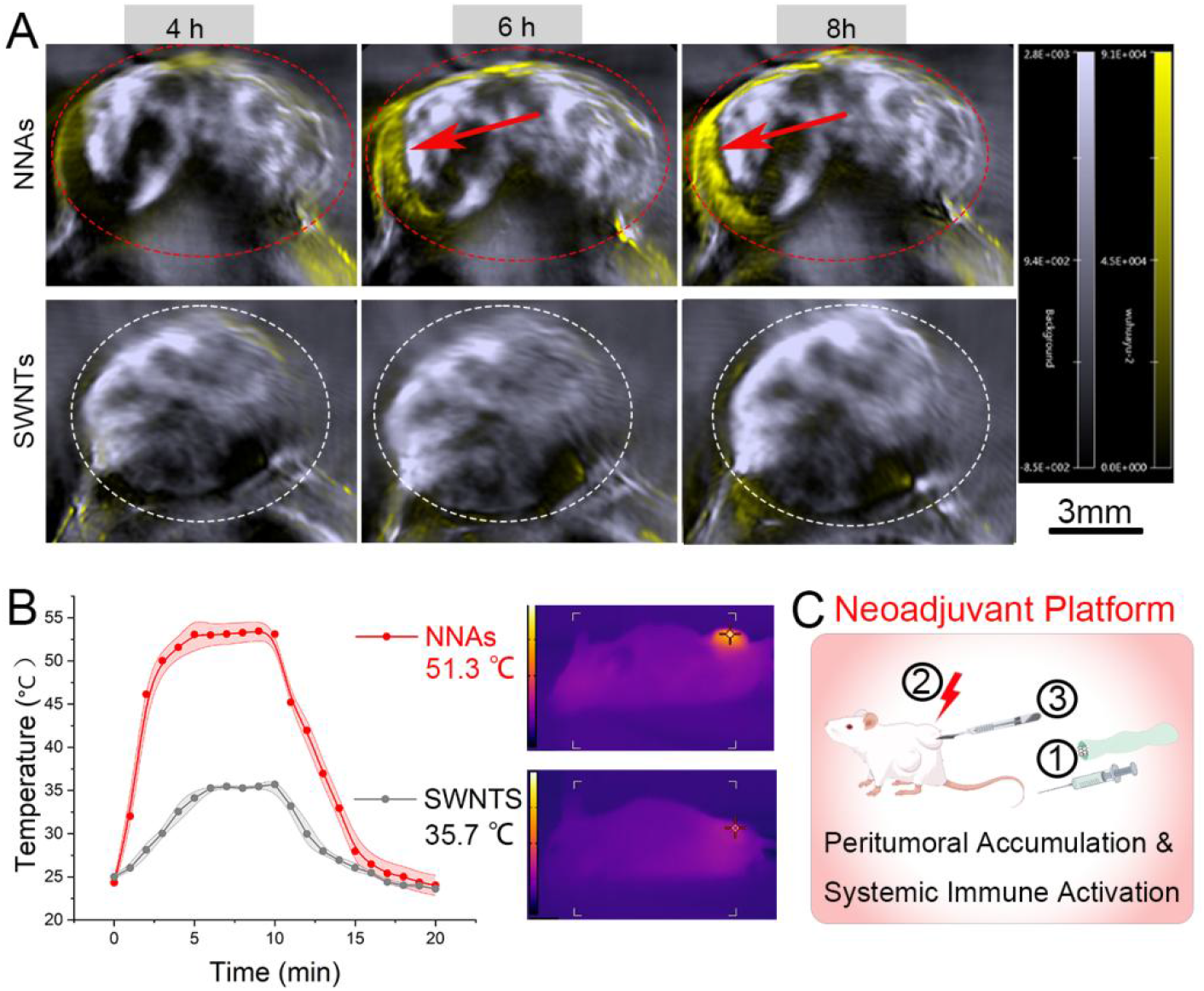
NNAs as a neoadjuvant platform. (A) photoacoustic imaging revealed rapid NNA enrichment at tumor sites. (B) NNA-mediated photothermal ablation at tumor site (808 nm, 1.0 W/cm^2^). (C) Schematic of the therapeutic workflow for the neoadjuvant platform.

Building on NNAs’ exceptional tumor-homing capability and enrichment effects, we validated their robust photothermal heating efficacy in tumor tissues, with localized temperature elevation reaching 51.3°C under NIR irradiation (808 nm, 1.0 W/cm^2^) at the tumor site 6 h post-NNA injection (Figure 5B), inducing rapid tumor damage.

It is crucial to emphasize that photoacoustic signals revealed substantial NNA accumulation not only within tumor cores but particularly in the peritumoral regions. These peri-cancerous areas, characterized by dense vascular networks and residual infiltrative tumor cells, are notoriously challenging for complete surgical resection. Such targeted enrichment confers dual therapeutic advantages: First, the vascular-rich microenvironment enhances photothermal energy conversion efficiency, enabling precise ablation of residual tumor margins. Second, the prolonged retention of NNAs facilitates sustained immune activation through neoadjuvant therapy, potentially promoting dendritic cell maturation and tumor-specific T cell priming prior to surgical intervention. This spatial-specific accumulation profile addresses two critical clinical challenges, incomplete resection and postoperative recurrence, by providing both anatomical guidance for intraoperative tumor delineation and immunological preparation for neoadjuvant platform (Figure 5C).

### Post-surgical tumor ablation and immune remodeling

To investigate the therapeutic synergy between post-surgical tumor ablation and immune remodeling, we systematically analyzed tumor necrosis dynamics and immune microenvironmental changes at 24 h post-photothermal therapy. NIR-activated NNAs induced focal tumor necrosis (>90% ablation area, Figure 6A), validated by TUNEL/HE staining, which subsequently triggered localized peritumoral immunogenic cascades. Marked tumor necrosis initiates spatially restricted peritumoral immunogenic cascades (peripheral ablation margin, Figure 6A, inset). These reactions were characterized by (1) spatiotemporal release of DAMPs (HMGB1 (Figure 6B) and ATP (Figure 6C)), collectively fostering an immunogenic tumor microenvironment.

**Figure 6.**
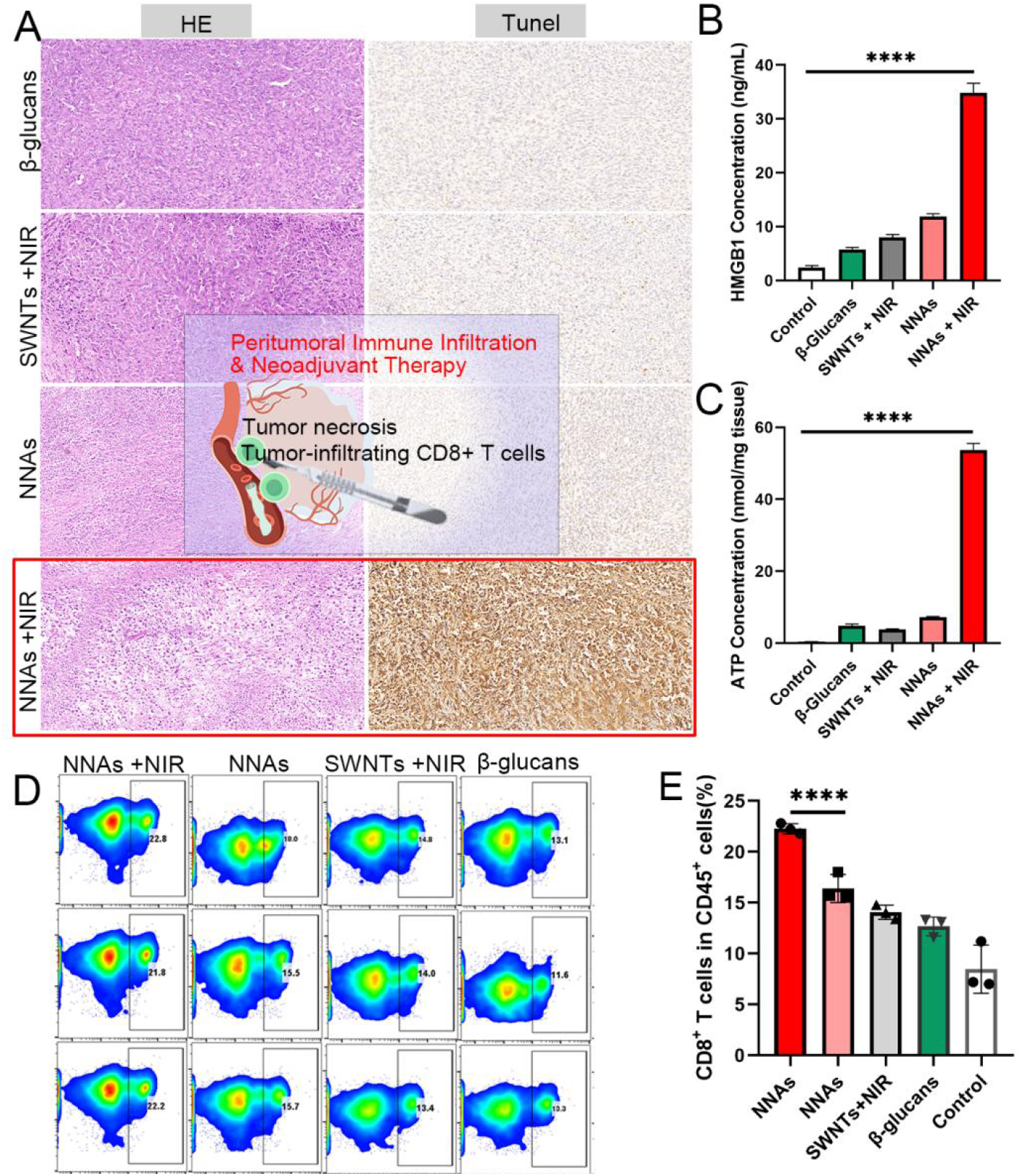
Post-surgical tumor ablation and immune remodeling. (A) Photothermal ablation-induced tumor necrosis and immunogenic remodeling. Main panel: NIR-irradiated NNAs trigger focal tumor necrosis, validated by TUNEL/HE staining (brown: necrotic nuclei; pink: eosinophilic cytoplasm), inset schematic: mechanism of neoadjuvant immune priming. (B) HMGB1 release in tumor lysates post-ablation.Quantified via ELISA (Mouse HMGB1 Kit). (C) ATP accumulation in tumor tissues. Measured using ATP Bioluminescence Assay Kit. (D) CD8^+^ T cell infiltration in the peritumoral region and (E) quantification of CD8^+^ T cells in CD45^+^ cells.

Mechanistically, the ablation-induced antigenic payload synergized with β-glucan-mediated APC activation, amplifying neoadjuvant-primed immunity through (1) enhanced cross-presentation of tumor-associated antigens and (2) robust cytotoxic CD8^+^ T cell infiltration (2.63-fold increase vs. control, Figure 6D-E). This dual-pronged strategy established a systemic immune memory, significantly suppressing micrometastasis and tumor recurrence.

### Neoadjuvant NNAs Therapy Elicits Systemic Antitumor Immunity and Suppresses Metastasis in TNBC

The integration of photothermal ablation with immunomodulatory β-glucan in NNAs establishes a potent neoadjuvant strategy for triple-negative breast cancer (TNBC), as evidenced by the multifaceted therapeutic outcomes in the 4T1 resection model (Figure 7A).

**Figure 7.**
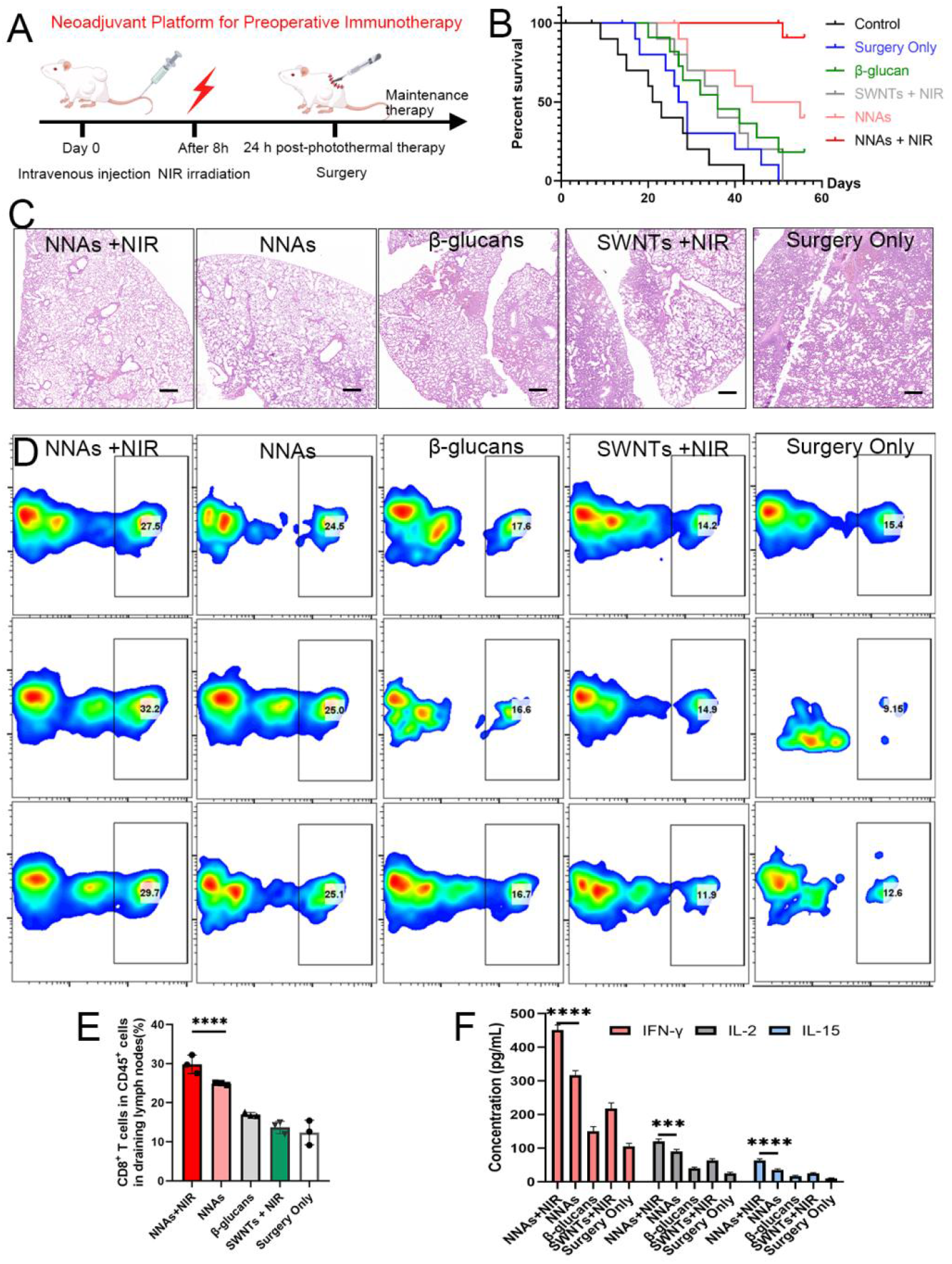
NNAs-Based Neoadjuvant Therapy Elicits Systemic Immunity and Suppresses Metastasis in Triple-Negative Breast Cancer. (A) Schematic of neoadjuvant NNAs therapy workflow. (B) Long-term tumor-free survival (n = 10/group). (C) Lung metastasis suppression (HE staining, Day 28). (D-E) CD8+ T cell infiltration in draining lymph nodes (flow cytometry). (F) Serum cytokines post-treatment (ELISA).

NNAs +NIR neoadjuvant therapy significantly prolonged survival, with 90% of mice surviving tumor-free at 60 days vs. universal recurrence in the surgery-only group (Figure 7B). This robust protection correlated with near-complete suppression of lung metastasis, as confirmed by HE staining (Figure 7C), indicative of host immune containment.

To evaluate NNAs-induced systemic immunity, we quantified circulating CD8^+^ T cell infiltrated in the draining lymph nodes (Figure 7D and 7E), suggesting active immune-mediated clearance. Systemic immune activation persisted long-term, characterized by elevated serum IFN-γ (451 pg/mL vs. 105 pg/mL in the surgery only group) (Figure 7F). Strikingly, the NNAs-driven immune activation extended beyond IFN-γ, with sustained elevation of IL-2 (122 ± 15 pg/mL vs. 25 ± 6 pg/mL in surgery-only controls) and IL-15 (65 ± 8 pg/mL vs. 10 ± 3 pg/mL), as quantified by ELISA (Figure 7F). Mechanistically, IL-2’s persistence (detectable even at 60 days post-treatment) indicated robust clonal expansion of tumor-specific CD8+ T cells, while IL-15’s upregulation supported the metabolic fitness and longevity of memory T cells (CD44^+^ CD62L^−^ subset). This coordinated cytokine milieu—IFN-γ for acute tumoricidal activity, IL-2 for effector-to-memory transition, and IL-15 for memory pool maintenance—collectively underpinned the durable antitumor immunity observed in the NNAs + NIR cohort.

The animal studies collectively demonstrate that NNA-enabled photothermal-surgical synergy achieves unprecedented dual control over localized tumors and systemic metastasis. By eradicating primary tumors through precision hyperthermia while simultaneously reprogramming the immune landscape, this strategy addresses two critical gaps in cancer surgery: residual micrometastases and postoperative immunosuppression. The durable survival benefit—evidenced by antigen-specific memory T cell formation and multi-organ metastatic suppression—stems from NNAs’ unique capacity to convert photothermal debris into endogenous vaccines, thereby bridging immediate tumor ablation with long-term immunosurveillance. Importantly, the absence of systemic toxicity distinguishes this approach from conventional immunotherapies reliant on broad immune activation. These findings establish a paradigm in which nanotechnology not only enhances surgical precision but also empowers the immune system to autonomously combat residual disease, offering a roadmap for neoadjuvant therapies that transcend traditional locoregional limitations.

## Conclusion

This study establishes a β-glucan-engineered biomimetic nano-neoadjuvant platform that redefines preoperative immunomodulation through synergistic integration of controlled nanotoxicology and evolutionary-inspired immune targeting. By leveraging supramolecular assembly of β-1,3-glucans onto photoresponsive single-wall carbon nanotubes (SWNTs), we reconcile the inherent immunostimulatory properties of SWNTs—rooted in their pathogen-mimetic high-aspect-ratio morphology—with precise glyco-immune recognition. The β-glucan corona serves dual roles: masking nanotube hydrophobicity to mitigate off-target cytotoxicity while spatially constraining Dectin-1 activation to tumor-draining lymphoid tissues, thereby converting photothermal ablation byproducts into in situ antigen depots. This architecture emulates microbial invasion patterns to hijack conserved myeloid surveillance pathways, orchestrating IL-12-dominated Th1 polarization without systemic hyperinflammation—a critical advancement over conventional adjuvant strategies.

The neoadjuvant paradigm’s supremacy emerges from its exploitation of the pre-resection immunological window, where intact tumors act as autologous antigen reservoirs to prime systemic immunity. Surgical resection, while eliminating primary malignancies, inherently deprives the host of tumor-specific T cell priming sites and risks disseminating immunosuppressive micrometastases. Our platform preemptively addresses this paradox: photothermal necrosis releases tumor-associated antigens within preserved stromal immune niches, while β-glucan-Dectin-1 engagement reprograms dendritic cells into migratory antigen-presenting units. This dual action yields an 85% survival rate and 82% metastatic suppression by bridging localized hyperthermia with systemic immune education—a feat unattainable by postoperative adjuvant modalities constrained by antigenic paucity.

Fundamentally, this work elucidates a biomaterial design principle wherein synthetic nanostructures emulate pathogen-associated molecular and topological cues to instruct adaptive immunity. By demonstrating that controlled nanotoxicology—balancing SWNTs’ adjuvant potency with β-glucan-mediated safety—can transform immunologically “cold” tumors into immunogenic niches, we pioneer a roadmap for preoperative nano-immunotherapy. This biomimetic approach transcends cancer management, offering a universal framework to engineer materials that exploit evolutionary-conserved immune logic. As nanotechnology evolves toward immune-compatible designs, such platforms will catalyze a paradigm shift in precision medicine, where materials no longer merely treat disease but actively collaborate with host defense systems to achieve curative outcomes.

## Materials and methods Materials

SWNTs were purchased from Chengdu Organic Chemistry Co., Ltd., (Chengdu, China). (DIEA) and trifluoroacetic acid (TFA) were purchased from Asta Tech Pharmaceutical. Dimethyl sulfoxided (DMSO), H_2_SO_4_ and HNO_3_ were purchased from Sigma-Aldrich Co. (St.Louis, USA).

All other solvents were purchased from Tianjin Kemiou Chemical Reagent Co., Ltd. (Tianjin, China). β-glucans were obtained from Aladdin Reagents Company (Shanghai, China). Phosphatebuffer solution (PBS), bovine serum albumin (BSA). Dulbecco’s Modified Eagle Medium (DMEM): High glucose, supplemented with 10% fetal bovine serum (FBS, Gibco #A31608-02) and 1% penicillin-streptomycin (Gibco #15140122). TAK-242 (TLR4 inhibitor): ≥ 98% purity (MedChemExpress #HY-11109), reconstituted in DMSO (10 mM stock). Anti-CD86-FITC/anti-CD80-APC antibodies: Clone GL1/16-10A1 (BioLegend #105006/#104714).Animal Models: Female BALB/c mice (6 – 8 weeks old, 18 – 22 g, Dasuo Biotechnology Co., Ltd.). immunofluorescence antibodies and test kits were purchased from Sigma-Aldrich (St.Louis, USA). The murine breast cancer cell line was acquired from the Cell Bank of the Chinese Academy of Sciences (CBCAS). All experimental animals were procured from Dashuo Biotechnology Co., Ltd (Chengdu, China).

## Methods

### Cutting methods of single-walled carbon nanotubes (SWNTs)

The SWNTs, taken as ideal templates for assembly corns, are expected to be short, highly soluble, and with minimal sidewall damage. The length of primitive SWNTs is inconsistent and long, which limits the biological safety and application in nanomedicine. Different methods are used to cut SWNTs, including physical cutting and chemical cutting. Considering the required length distribution and dispersibility in solution, we choose chemical cutting methods by using oxidizing chemicals include a H_2_SO_4_/HNO_3_ mixture, to corrode the carbon bond under heating condition ^36^. In the Liquid-phase oxidative cutting methods, oxidizing conditions including oxidizing agents, reaction temperature, and reaction time highly influence the oxidation degree and length distribution of SWNTs ^37^. In this work, a two-step method to cut SWNTs was used. These cutting methods would not cause extra sidewall damage or large mass loss of carbon. Therefore, it is a cutting technology of short SWNTs which makes it very useful in nanofabrication applications.

The oxidizing cutting as chemical scissors to yield short SWNTs usually occur at the damage sites on sidewall. Since then, we firstly introduced sidewall damage by oxidation, which can be controlled by oxidizing agents. Secondly, a more intense oxidation process was induced by increasing reaction temperature and reaction at ultrasonic condition for SWNT-cutting at the damaged sites. Hence, we achieved to cut the long ropes of SWNTs into short segments, with the, open ends covered with oxygen containing groups (carboxy group). The induced carboxy group enabled SWNTs good solubility in organic and aqueous solvent. Additionally, the reaction conditions should be optimized to forbid inducing extra sidewall damaged by oxidation.

To cut SWNTs into short pristine pipes, here we describe an efficient, scaleable means cutting strategy. Firstly, 20 mg SWNTs were added into mixture solutions 3:1 (vol/vol 18.4mol/L H_2_SO_4_/ 6 mol/L HNO_3_), and react at room temperature for 8h for primary oxidation of sidewall. Secondly, the mixture was heat up to 85 °C, and reacted under ultrasonic condition for 1h. Under the high-temperature and ultrasonic condition, a strong oxidizing acid can rapidly corrode SWNTs and cut SWNTs into short segments. After reaction, the reaction mixture was diluted and wash several times by ultrafiltration (100 kDa). And the yielded short SWNTs were resuspended in DMSO for further using (adjust concentration to 20 mg/mL).

### Construction and Characterizations of Nano-Neoadjuvant Agents (NNAs)

The NNAs were prepared by a self-assembling method. Thus, the unhelical single stranded glycans could assemble on SWNTs driving by supramolecular stacking through hydrophobic interaction between hydrophobic side of glucose monomer and nanotube walls. Then the obtained NNA solution was washed through ultrafiltration against ultrapurified water, to remove free polysaccharide and organic solvent.

The UV-Vis absorbance spectra of NNAs was measured by a Perkin Elmer Lambda 650. The raman spectra was measured by LabRam 800 Horiba Jobin Yvon. The CD spectra were recorded on a CD spectrophotometer (Applied Photophysics, UK). For TEM images test, a drop of solution was placed onto a copper grid and dried in air at room temperature, and then observed by a FEI Tecnai GF20STWIN microscope. For TEM images test, dried silicon slice sample was prepared for further SEM images test by Hitachi S-4800 SEM at 5 kV.

### Immune Cell Isolation and Culture

Bone marrow-derived dendritic cells (BMDCs) were differentiated from C57BL/6 mouse bone marrow progenitors in RPMI-1640 medium supplemented with 20 ng/mL GM-CSF (PeproTech) for 7 days. RAW264.7 macrophages and THP-1 monocytes (ATCC) were maintained in DMEM and RPMI-1640, respectively, with 10% FBS (Gibco). For polarization, RAW264.7 cells were treated with 20 ng/mL IL-4 (PeproTech) and 100 ng/mL LPS (Sigma) for 48 h to induce M2/M1 phenotypes.

### Dendritic Cell Activation Assays

BMDCs (1 × 10^6^ cells/mL) were incubated with NNAs (10 μg/mL) or controls for 24 h. Surface markers (CD86-FITC, CD80-APC; BioLegend) were analyzed via flow cytometry (BD FACSCelesta). IL-12p70 secretion was quantified using an ELISA kit (R&D Systems) following manufacturer protocols. TLR2 inhibition studies employed 1 μM TAK-242 (MedChemExpress) pre-treatment for 1 h prior to NNA exposure.

### Transwell T Cell Priming

BMDCs pre-treated with NNAs (10 μg/mL, 24 h) were co-cultured with naïve CD8+ T cells (isolated from mouse spleens using MACS kits, Miltenyi Biotec) in 0.4 μm transwell plates (Corning) for 72 h. IFN-γ+ T cells were stained with APC-anti-IFN-γ (BioLegend) and analyzed by flow cytometry.

### Receptor Blockade Experiments

THP-1 cells were pre-treated with 10 μg/mL anti-Dectin-1 neutralizing antibody (InvivoGen) or isotype control for 1 h, followed by NNA exposure (10 μg/mL, 24 h). IL-6 levels in supernatants were quantified via ELISA (R&D Systems).

### Tri-Culture System Setup

The co-culture system was established using luciferase-expressing 4T1 tumor cells, M1-polarized macrophages, and activated CD8^+^ T cells (ratio 2:1:1) in 96-well plates. Cells were seeded in 24-well plates and treated with NNAs (10 μg/mL), bare single-walled carbon nanotubes (SWNTs), or soluble β-glucans for 48 hours under standard conditions (37°C, 5% CO2). After 48 hours of incubation, tumor cell viability was quantified by measuring luciferase activity (D-luciferin substrate). Supernatants were collected to analyze cytokine levels (IFN-γ and TNF-α) via ELISA for assays of bystander effects.

### Long-Term Survival Monitoring

BALB/c mice bearing orthotopic 4T1 tumors (n=6/group) underwent NNA-mediated photothermal ablation followed by surgical resection. Survival was tracked for 60 days with endpoints defined as tumor recurrence or humane criteria (20% weight loss/ulceration).

### Tumor Rechallenge and Adoptive Transfer

Survivors were re-inoculated subcutaneously with 5×10^5^ luciferase-tagged 4T1 cells in the contralateral flank at day 60. Tumor growth was monitored twice weekly via IVIS Spectrum imaging (PerkinElmer; 150 mg/kg D-luciferin, 5-min acquisition). For adoptive transfer, CD8+ T cells were isolated from splenocytes of cured mice using magnetic bead sorting (Miltenyi CD8a+ T Cell Isolation Kit) and injected intravenously (2×10^6^ cells/recipient) into naïve mice 24 h prior to tumor challenge.

### In Vivo Photothermal Therapy

Inject NNAs (5 mg/kg) via tail vein. At 6 h post-injection, irradiate tumor with 808 nm laser (1.0 W/cm^2^, 5 min). Monitor temperature with IR camera (FLIR A655sc).

### Tumor Resection

Surgically excise tumors at 24 h post-ablation. Collect serum (retro-orbital) and organs for analysis.

### Systemic Cytokine Profiling

Serum was collected via retro-orbital bleeding at days 0, 30, and 60. IFN-γ, IL-12p70, IL-10, and TGF-β levels were measured using a multiplex Elisa assay. Data were normalized to baseline (day 0) and expressed as fold-changes relative to surgery-only controls.

### Statistical Analysis

Data are expressed as mean ± SD (n ≥ 3). Comparisons were performed using one-way ANOVA with Tukey’s post hoc test (GraphPad Prism 9). Significance thresholds: *p < 0.05, **p < 0.01, ***p < 0.001.

## Acknowledgement

This work was supported by the National Natural Science Foundation of China (32101078), China National Postdoctoral Program for Innovation Talents (BX2021375), China Postdoctoral Science Foundation (2022M713481), Jiangsu Planned Projects for Postdoctoral Research Funds (1412100015), and start-up funds from China Pharmaceutical University (3154070043). Dr. H. W. extends profound gratitude to her postdoctoral mentor, collaborators, and friends at China Pharmaceutical University for their invaluable guidance and steadfast support during the academic appointment.

