## Supplementary Figure for "Supramolecular Biomimetic Topology: Macrophage-Engaged Neoadjuvant Platform for Preoperative Immunotherapy": Supporting information.pdf

**Biomimetic Neoadjuvant Platform: Controlled Nanotoxicology as a Novel Approach for Preoperative Immunomodulation**

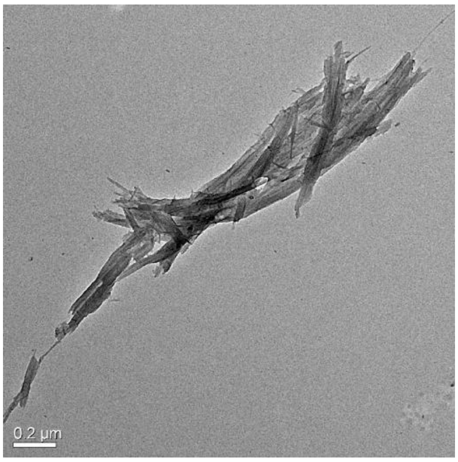

Supplementary Fig. 1. TEM images of non-cut SWNTs exhibited pronounced aggregation

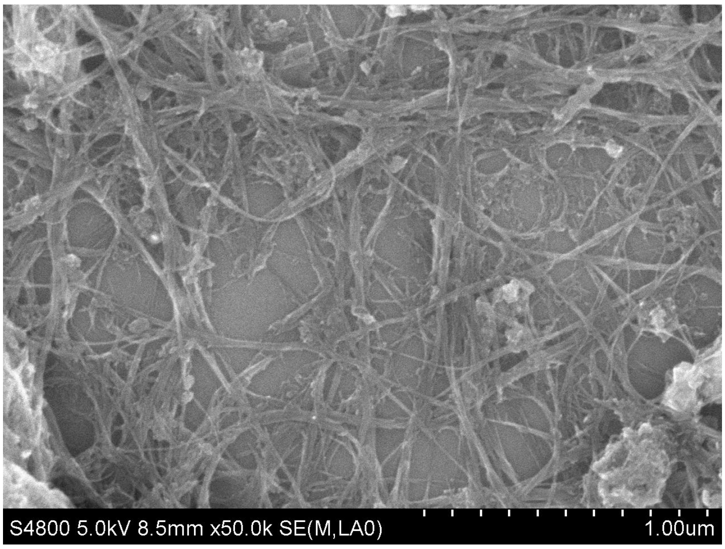

Supplementary Fig. 2. SEM images of non-cut SWNTs exhibited pronounced aggregation

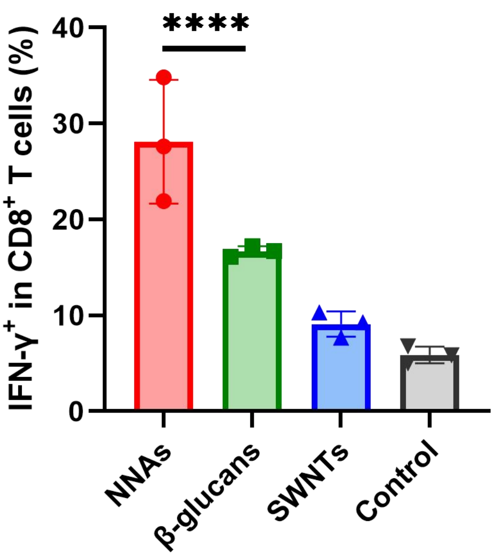


Supplementary Fig. 3. IFN-$\gamma^+$ CD8$^+$ T cells
